## Supplementary material for "Molecular mechanisms of the CdnG-Cap5 antiphage defense system employing 3’,2’-cGAMP as the second messenger": Fig. S1-S9, Table S1

Supplementary Information

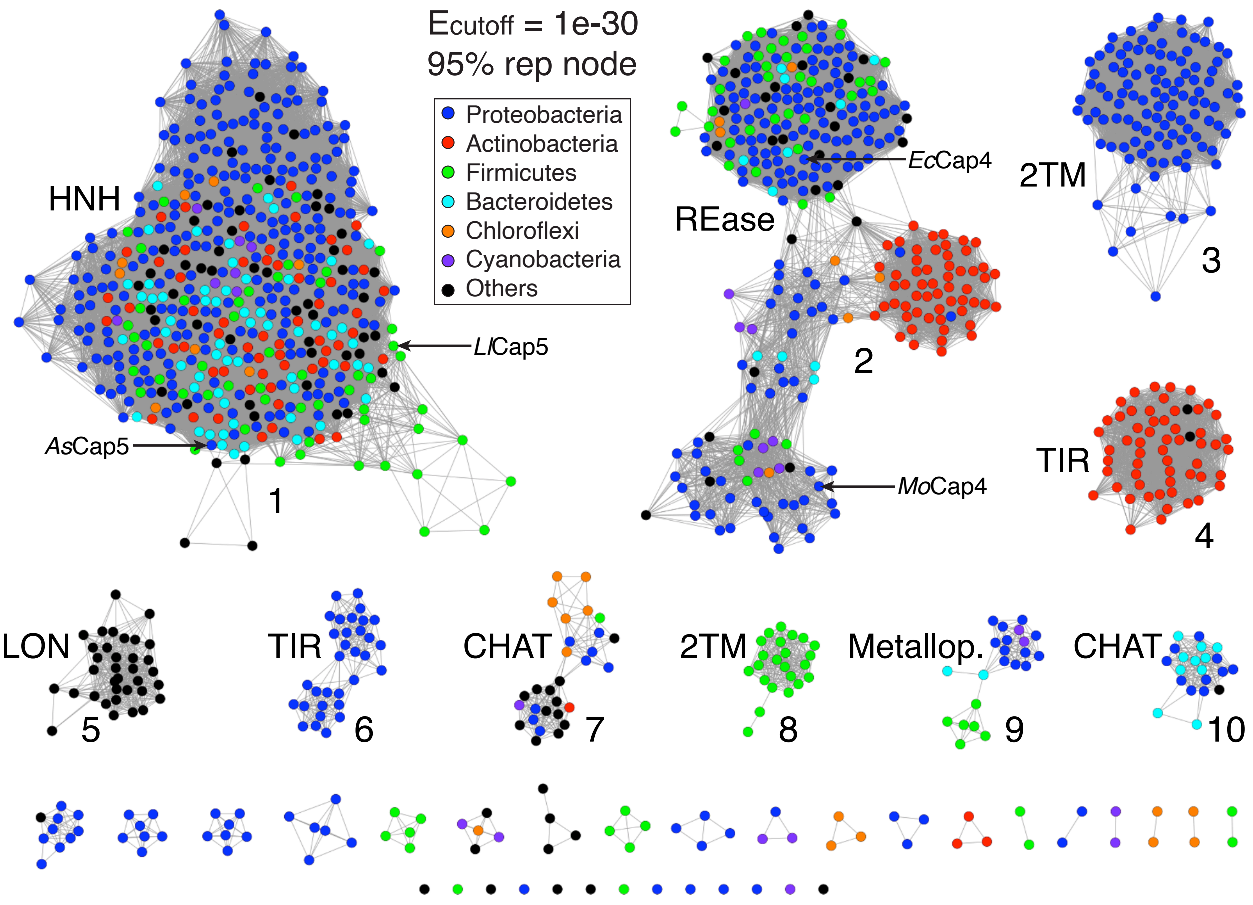


**Supplementary Figure 1. Bioinformatic analysis of SAVED domain-containing proteins via Sequence Similarity Network (SSN).** The analysis was based on sequences of PF18145 on the database of UniProt 2020_04 and InterPro 81. Each node (colored circle) represents a collection of SAVED proteins sharing >95% sequence identity (95% rep node). An edge (gray line) connects two nodes if the E-value measuring their sequence similarities is smaller than the cutoff value (1e-30 for this network). Only top 10 most abundant clusters are labeled, including abbreviations of effector domains associated with SAVED domain. The nodes representing two Cap5 of current study are marked with arrows. Also marked are two Cap4 proteins recently studied by Kranzusch and coworkers.

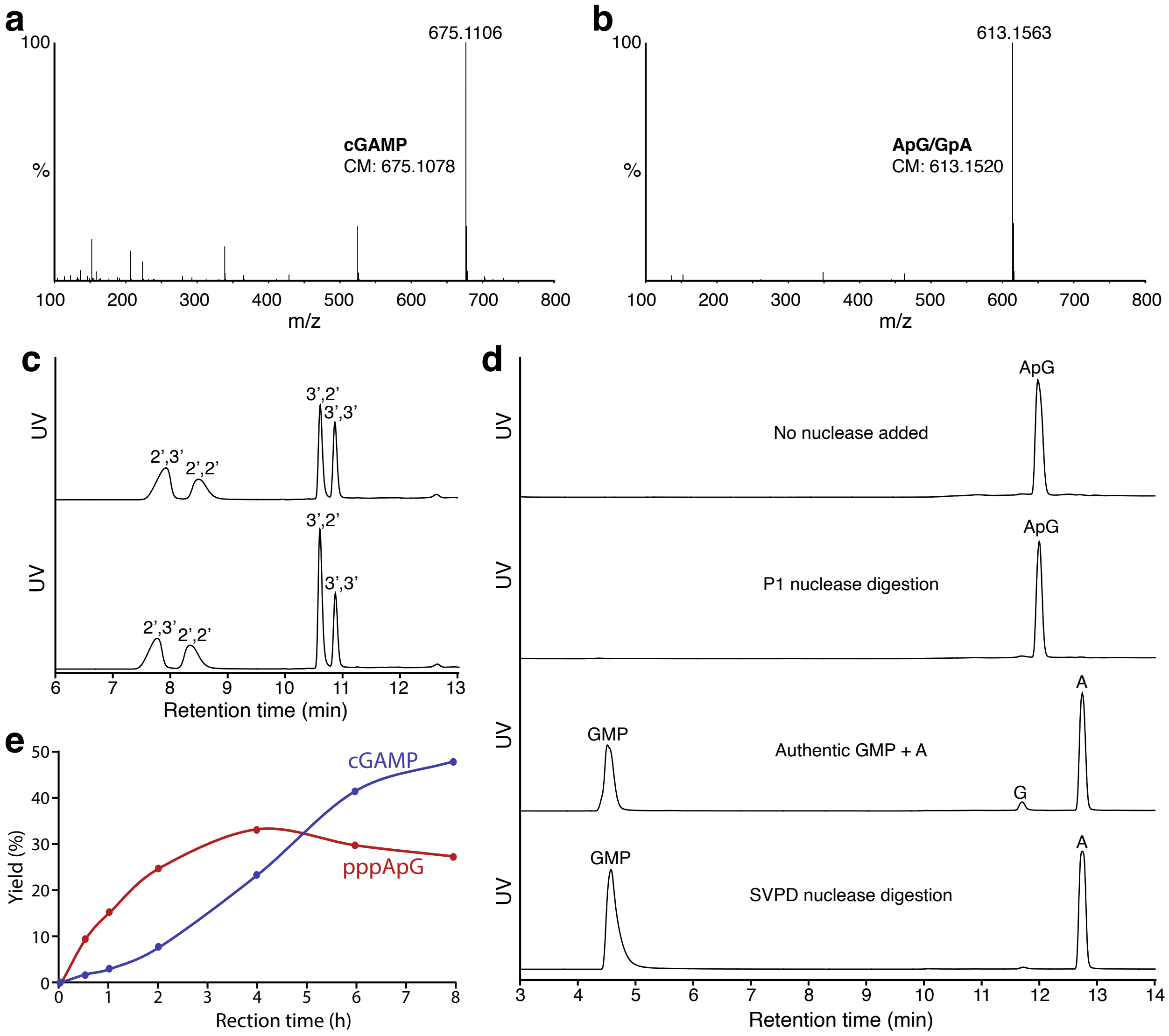


**Supplementary Figure 2. Characterization of chemical structures of two compounds synthesized by *As*CdnG. a**,**b** High-resolution MS of the two compounds produced by *As*CdnG shown in Fig. 1b. CM, calculated m/z. **c** UPLC analysis of four commercially purchased cGAMP (top panel) as well as their co-injection with cGAMP synthesized by *As*CdnG (bottom panel). **d** UPLC analysis of the digested products of ApG/GpA by P1 nuclease (top panel) and snake venom phosphodiesterase (SVPD) nuclease (bottom panel). P1 nuclease only cleaves 3',5'-phosphate linkage, whereas SVPD is able to cleave both 3',5'- and 2',5'-phosphate linkages. **e** Time course of the reaction carried out by *As*CdnG.

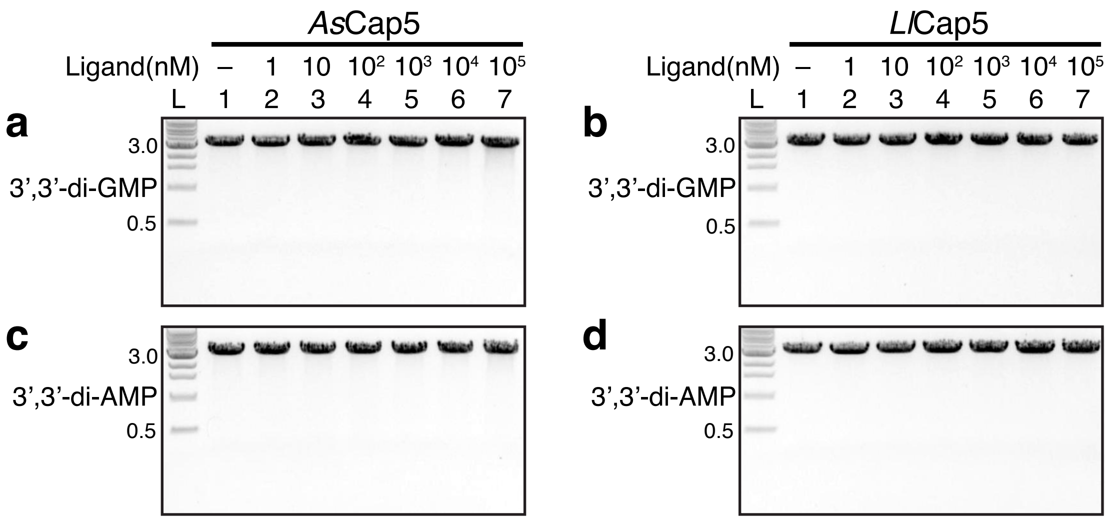


**Supplementary Figure 3. In vitro DNA degradation by *As*Cap5 and *Ll*Cap5 in the presence of 3',3'-c-di-GMP and 3',3'-c-di-AMP.** The experimental conditions were the same as the ones shown in Fig. 2.

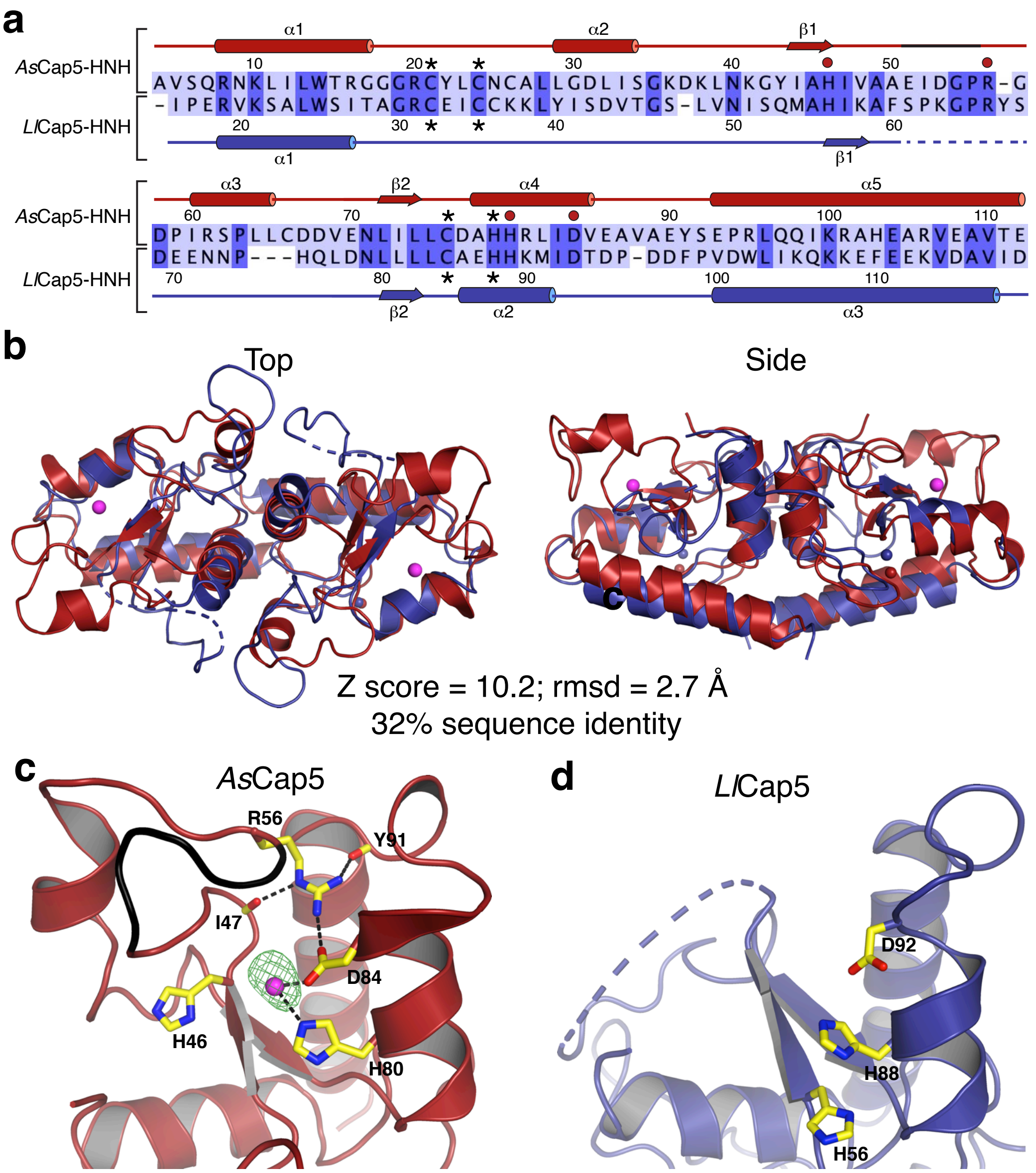


**Supplementary Figure 4. Additional structural analysis of Cap5-HNH. a** Mapping secondary structures of *As*Cap5-HNH (top) and *Ll*Cap5-HNH (bottom) onto their aligned sequences. Part of the loop between β1 and α3 in *As*Cap5-HNH having steric clash with the modeled DNA•RNA hybrid is colored black. The disordered residues in *Ll*Cap5-HNH are depicted with dashed lines. Zinc finger residues are marked with asterisks. Important residues near the active site of *As*Cap5-HNH are marked with red circles. **b** Structural alignments of *As*Cap5-HNH and *Ll*Cap5-HNH homodimers. The structures are depicted in cartoon, with *As*Cap5-HNH and *Ll*Cap5-HNH colored red and blue, respectively. **c** Structure of *As*Cap5-HNH near the active site. The $Mg^{2+}$ ion is covered with simulated-annealing $F_o$-$F_c$ omit map contoured at 3 σ. H46 activates a water molecule to cleave DNA. Hydrogen bonding network of the side chain of R56 is likely to be responsible for the conformation of the loop that has steric clash with the modeled DNA•RNA hybrid. **d** Structure of *Ll*Cap5-HNH with the same region as *As*Cap5-HNH shown in **c**, indicating that *Ll*Cap5-HNH is significantly less structured near the active site.

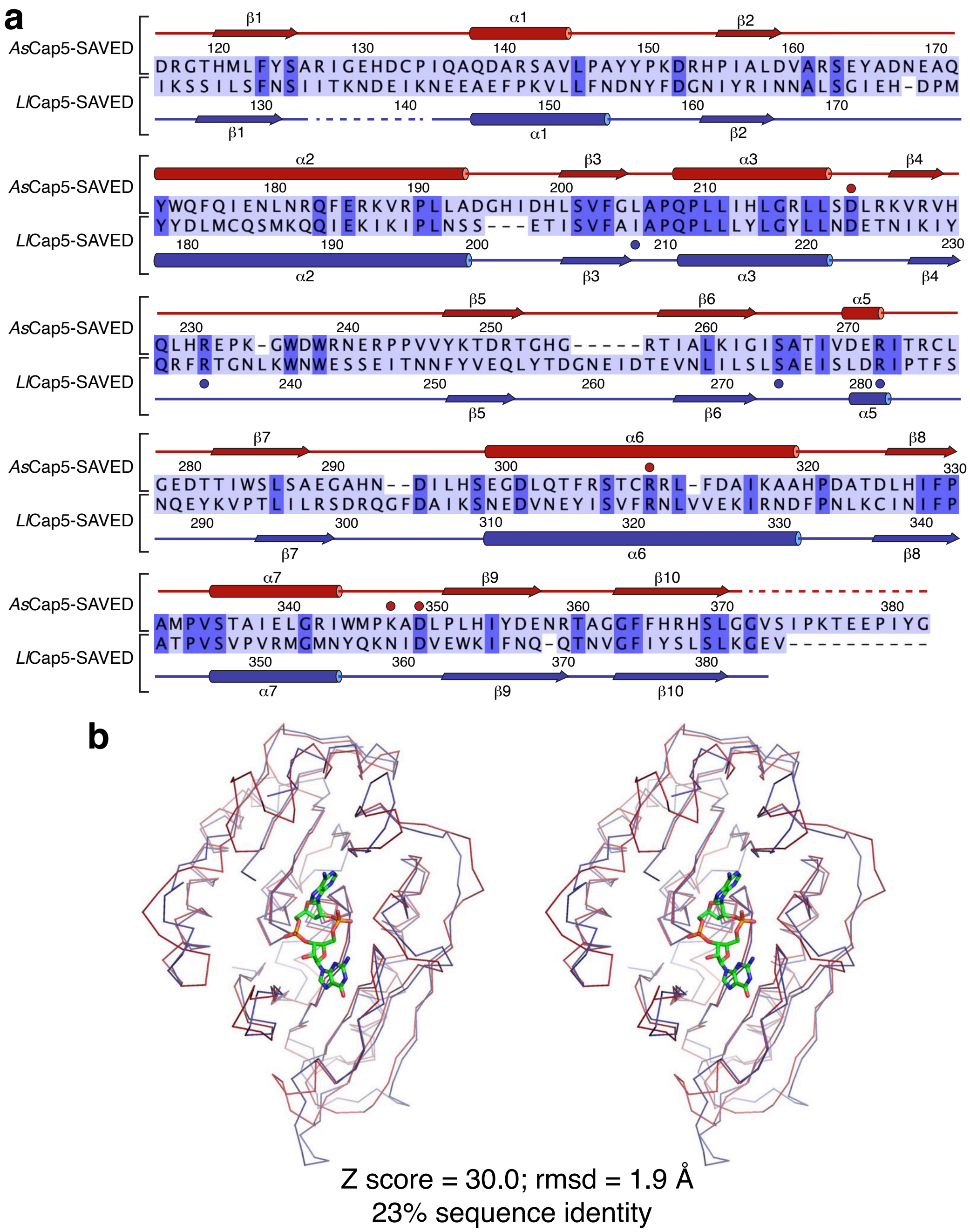


**Supplementary Figure 5. Additional structural analysis of the SAVED domains of Cap5. a** Mapping secondary structures of *As*Cap5-SAVED (top) and *Ll*Cap5-SAVED (bottom) onto their aligned sequences. The two SAVED domains only share 23% sequence identity. Residues of *Ll*Cap5-SAVED involved in recognition of 3',2'-cGAMP are marked with blue circles. Residues of *As*Cap5-SAVED selected for mutagenesis are marked with red circles. **b** Stereoview of structural superposition of *As*Cap5-SAVED (red) and *Ll*Cap5-SAVED (blue). The structures are depicted in ribbon, and the bound 3',2'-cGAMP is in stick.

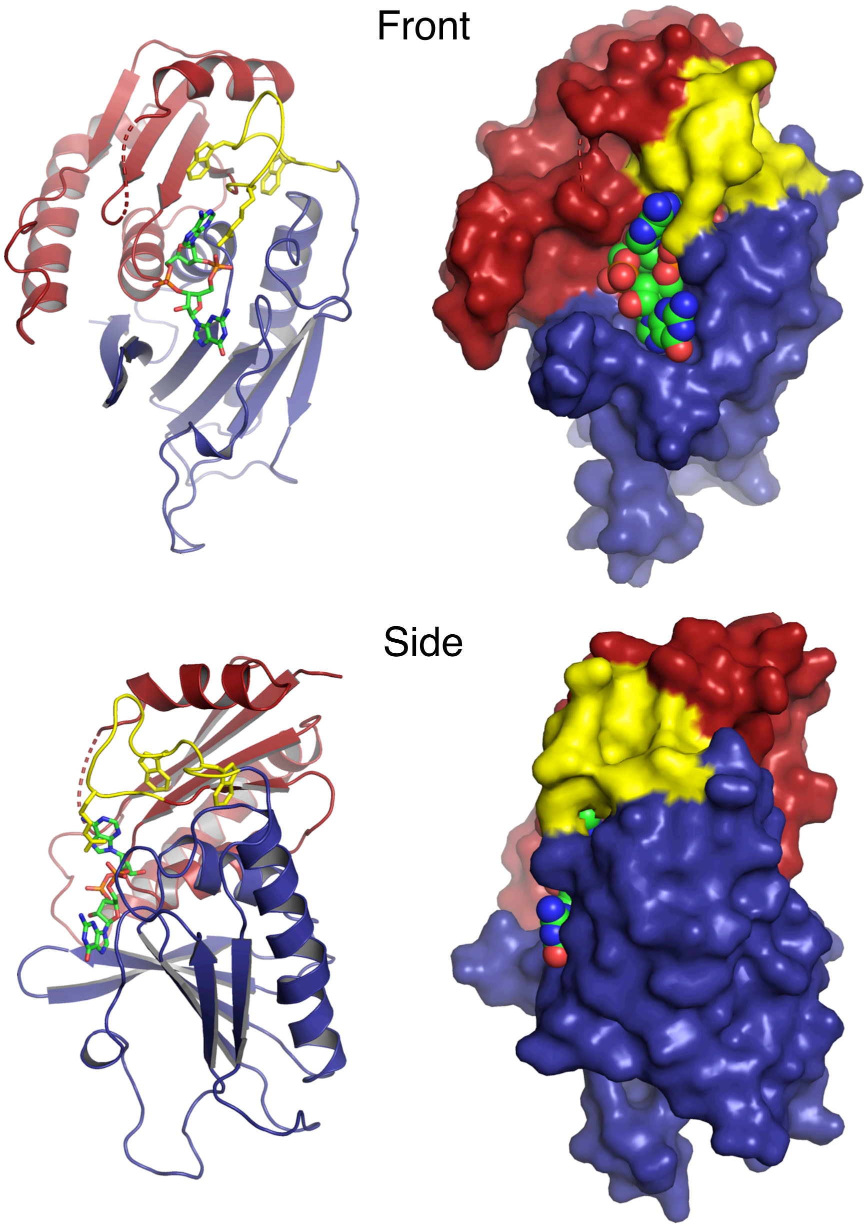


**Supplementary Figure 6. Structural analysis of *Ll*Cap5-SAVED-3',2'-cGAMP complex in the context of possible evolutionary origin of SAVED domain.** The structures are depicted in Cartoon (left) and Surface (right), and oriented the same as the ones shown in Fig. 4a,b. The first CARF-like motif is colored red, and the second one is colored blue. The linker connecting these two CARF-like motifs is colored yellow. The critical residue R234, as well as the conserved WxW motif, are in the linker region and highlighted in stick. Interactions of WxW motif with its surroundings are likely to play a role of orienting R234 for its interaction with 3',2'-cGAMP.

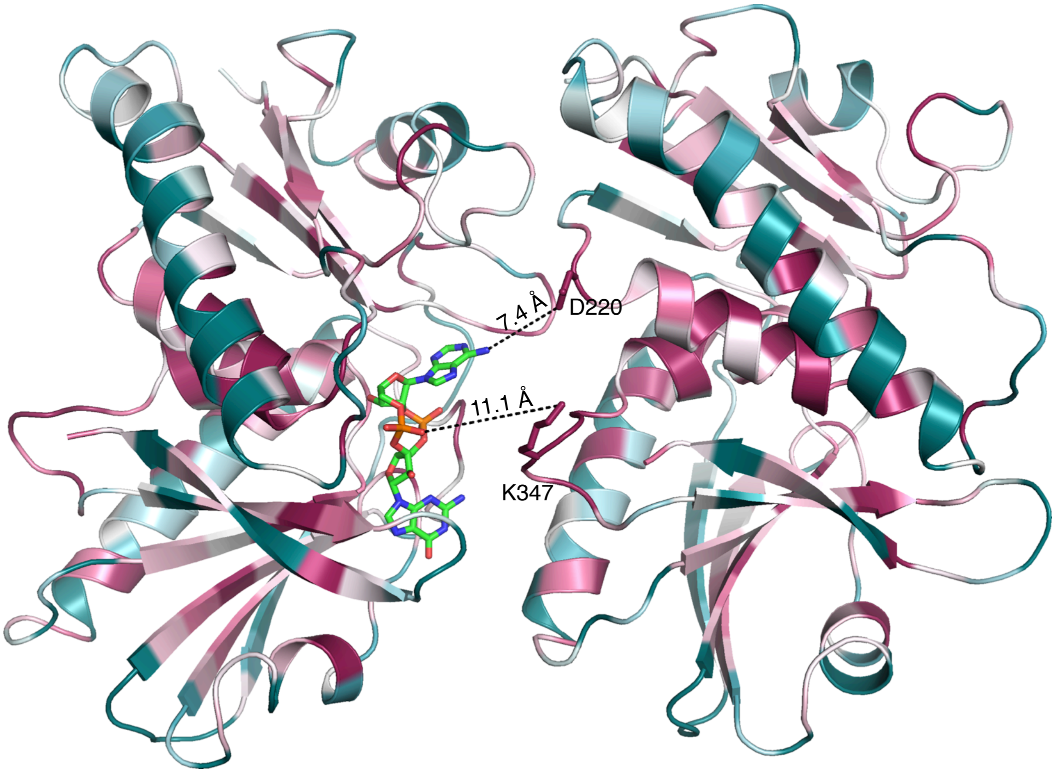


**Supplementary Figure 7. Cartoon depiction of the structure of *As*Cap5 SAVED dimer.** The structure is the same as the one shown in Fig. 5a with the addition of the modeled 3’,2’-cGAMP. Two conserved residues, D220 and K347, located at the tip of two extruding loops, are in stick. These two residues are proposed to interact with 3’,2’-cGAMP. We speculate that the side chain of D220 might interact with N6 of the AMP moiety, which is 7.4 Å away. We further speculate that the side chain of K347 might interact with the 3’,5’-phosphate group of the AMP moiety, which is 11.1 Å away based on the current structure. An optimal rotamer of K347 would reduce the distance to 7.2 Å. Therefore, if 3’,2’-cGAMP indeed induce two SAVED domains to form closed homodimer, the relative movement would be approximately 5 Å from the structure shown here.

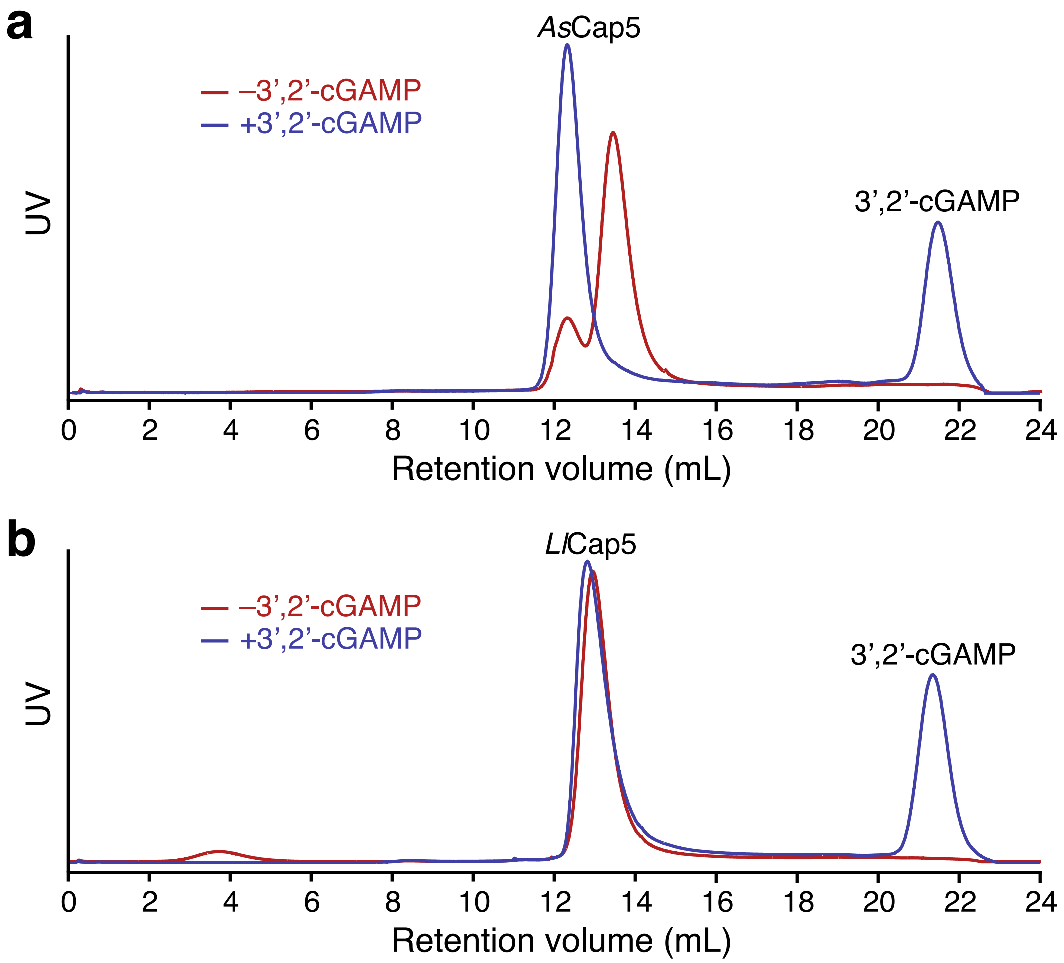


**Supplementary Figure 8. *As*Cap5 and *Ll*Cap5 do not form oligomers regardless of whether 3',2'-cGAMP is present.** Purified recombinant *As*Cap5 and *Ll*Cap5 were analyzed by analytical size exclusion chromatography in the absence (red curve) and presence (blue curve) of 3',2'-cGAMP. The peaks corresponding to the excess 3',2'-cGAMP were only observed in the samples with 3',2'-cGAMP (blue curves). 3',2'-cGAMP induced the shift of the retention time of *As*Cap5 (**a**). On the other hand, 3',2'-cGAMP had no effect on *Ll*Cap5 (**b**). Because both the size and the shape of a protein affect its retention volume of a gel filtration column, the results shown here are difficult to interpret. It is possible that the two peaks shown in **a** represent two different conformers of *As*Cap5 homodimer, and addition of 3',2'-cGAMP shifts the relative populations of these two conformers. One possible interpretation of the results shown in **b** is that the exchange of conformers of *Ll*Cap5 is too fast to be resolved by size exclusion chromatography. However, one conclusion is certain with these results: both *As*Cap5 and *Ll*Cap5 do not form oligomers in the presence of 3',2'-cGAMP under the conditions these analyses were carried out. In addition, the fractions collected from both *As*Cap5 and *Ll*Cap5 samples with 3',2'-cGAMP were active based on DNA degradation assays, demonstrating that 3',2'-cGAMP did not dissociate from the proteins over the course of size exclusion chromatography.

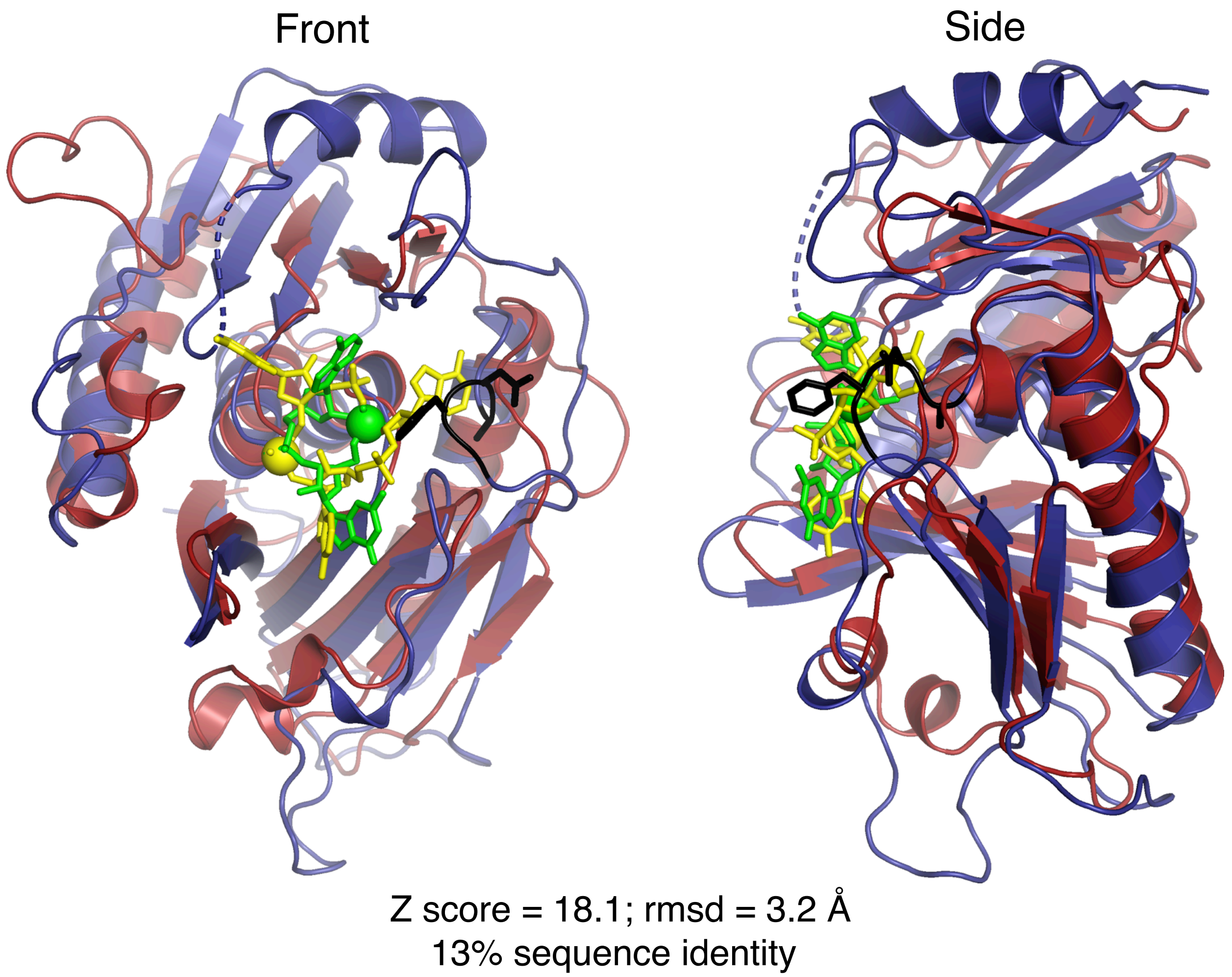


**Supplementary Figure 9. Structural comparison of *Ll*Cap5-SAVED to *Ab*Cap4-SAVED.** Two superpositioned structures are depicted in cartoon, with *Ll*Cap5-SAVED colored blue and *Ab*Cap4-SAVED colored red. The structures are oriented the same as the ones shown in Fig. 4a and Supplementary Figure 6. The bound ligands are depicted in stick, with 3',2'-cGAMP colored green and 2',3',3'-cAAA colored yellow. The phosphate atoms of the 2',5'-phosphate linkages are highlighted in spheres. Part of the loop of *Ll*Cap5-SAVED having steric clash with the third nucleotide of 2',3',3'-cAAA is colored black and depicted in stick.

### Supplementary Table 1. Data collection and model statistics

| | ***As*Cap5** | ***Ll*Cap5** | ***Ll*Cap5-SAVED + 3',2'-cGAMP** |
|---|---|---|---|
| PDB ID | 7RWK | 7RWM | 7RWS |
| **Data Collection** | | | |
| Space Group | $P2_12_12_1$ | $P4_3$ | $C222_1$ |
| Cell dimensions<br>a, b, c (Å)<br>$\alpha, \beta, \gamma$ (°) | <br>69.3 96.9 131.8<br>90.0, 90.0, 90.0 | <br>180.8 180.8 93.5<br>90.0, 90.0, 90.0 | <br>68.7 126.7 83.4<br>90.0, 90.0, 90.0 |
| Resolution (Å) | 50.0-2.4 | 50.0-3.4 | 50.0-1.8 |
| $R_{merge}$ (%) | 6.9 | 6.1 | 5.2 |
| $I/\sigma$ | 10.3 | 9.2 | 16.4 |
| Completeness (%) | 99.8 | 100 | 99.7 |
| Redundancy | 3.8 | 7.6 | 5.7 |
| **Refinement** | | | |
| Resolution (Å) | 39.7-2.4 (2.48-2.4) | 43.8-3.4 (3.52-3.4) | 33.0-1.8 (1.86-1.8) |
| No. reflections | 35,452 (3,305) | 41,617 (4,116) | 33,873 (3,297) |
| $R_{work}/R_{free}$ (%) | 19.3/23.5 (24.4/27.2) | 24.9/29.5 (27.5/36.6) | 16.6/19.3 (16.8/21.5) |
| No. atoms<br>Protein<br>Ligand/ion<br>Solvent | 6,223<br>5,858<br>4<br>361 | 11,431<br>11,427<br>4<br>0 | 2,451<br>2,073<br>45<br>333 |
| Average B factors (Å$^2$)<br>Protein<br>Ligand/ion<br>Water | 39.6<br>39.6<br>34.1<br>40.1 | 87.4<br>87.4<br>83.4<br>N/A | 21.9<br>20.1<br>15.2<br>33.8 |
| R.m.s. deviations<br>Bond lengths (Å)<br>Bond angles (°) | <br>0.010<br>1.55 | <br>0.040<br>0.88 | <br>0.010<br>1.86 |
| Ramachandran statistics (%)<br>Favored<br>Allowed<br>Outliers | <br>95.5<br>4.5<br>0.0 | <br>89.4<br>9.2<br>1.4 | <br>96.4<br>3.2<br>0.4 |

*Values in parenthesis are for highest resolution shell.
